# Repair of APOBEC3G-mutated retroviral DNA *in vivo* is facilitated by the host enzyme uracil DNA glycosylase 2

**DOI:** 10.1101/2021.08.25.457690

**Authors:** Karen Salas Briceno, Susan R. Ross

## Abstract

Apolipoprotein B mRNA Editing Enzyme Catalytic Subunit 3 (APOBEC3) proteins are critical for the control of infection by retroviruses. These proteins deaminate cytidines in negative strand DNA during reverse transcription, leading to G to A changes in coding strands. Uracil DNA glycosylase (UNG) is a host enzyme that excises uracils in genomic DNA, which the base excision repair machinery then repairs. Whether UNG removes uracils found in retroviral DNA after APOBEC3-mediated mutation is not clear, and whether this occurs *in vivo* has not been demonstrated. To determine if UNG plays a role in the repair of retroviral DNA, we used APOBEC3G (A3G) transgenic mice which we showed previously had extensive deamination of murine leukemia virus (MLV) proviruses. The A3G transgene was crossed onto an UNG and mouse APOBEC3 knockout background (UNG-/-APO-/-) and the mice were infected with MLV. We found that virus infection levels were decreased in A3G UNG-/-APO-/- compared to A3G APO-/- mice. Deep sequencing of the proviruses showed that there were significantly higher levels of G-to-A mutations in proviral DNA from A3G transgenic UNG-/-APO-/- than A3G transgenic APO-/- mice, suggesting that UNG plays a role in the repair of uracil-containing proviruses. In in vitro studies, we found that cytoplasmic viral DNA deaminated by APOBEC3G was uracilated. In the absence of UNG, the uracil-containing proviruses integrated at higher levels into the genome than did those made in the presence of UNG. Thus, UNG also functions in the nucleus prior to integration by nicking uracil-containing viral DNA, thereby blocking integration. These data show that UNG plays a critical role in the repair of the damage inflicted by APOBEC3 deamination of reverse-transcribed DNA.

**Importance:** While APOBEC3-mediated mutation of retroviruses is well-established, what role the host base excision repair enzymes play in correcting these mutations is not clear. This question is especially difficult to address *in vivo*. Here, we use a transgenic mouse developed by our lab that expresses human APOBEC3G and also lacks the endogenous uracil DNA glycosylase (*Ung*) gene, and show that UNG removes uracils introduced by this cytidine deaminase in MLV reverse transcripts, thereby reducing G-to-A mutations in proviruses. Furthermore, our data suggest that UNG removes uracils at two stages in infection – in unintegrated nuclear viral reverse transcribed DNA, resulting in its degradation and second, in integrated proviruses, resulting in their repair. These data suggest that retroviruses damaged by host cytidine deaminases take advantage of the host DNA repair system to overcome this damage.

## Introduction

Organisms adapt to infectious agents by developing protective responses and conversely, these agents develop adaptive countermeasures to these responses. Host defenses against infectious agents include various mechanisms of innate and adaptive immunity. One such family of host factors, apolipoprotein B mRNA editing enzyme, catalytic subunit 3 (APOBEC3) proteins belongs to a larger gene family encoding DNA and RNA editing enzymes characterized by the presence of at least one cytidine deaminase (CDA) domain (1). This family includes the Activation-Induced Cytidine Deaminase (AID) protein which is responsible for class-switch recombination and somatic hypermutation of the B cell receptor locus during germinal center development in lymph nodes, thereby contributing to antibody diversity (2). Cytidine deaminases, as well as other mutagens such as UV light, cause C-to-U changes in genomic DNA, which are then read as thymidines by DNA polymerase. (3). As such, the base excision repair (BER) machinery, including the nuclear form of Uracil DNA Glycosylase (UNG), removes uracils from DNA and in conjunction with other BER proteins, restores the original sequence, although since this latter process is error-prone, also causes mutations (4). There are two UNG splice variants; the mitochondrial form, UNG1, performs a similar role in this compartment (5). If UNG2, hereafter referred to as UNG, is not present, the uracils are read as thymines by DNA polymerase II and G-to-A transitions in the opposite strand occur (6). While UNG works on both double-strand (ds) with mismatches, its preferred template is single-strand DNA (7).

When packaged into retroviral virions, APOBEC3 proteins inhibit infection in target cells by deaminating deoxycytidine residues on minus strand DNA, causing G-to-A mutations in newly synthesized retrovirus coding strand DNA (8, 9). Deamination leads to degradation of reversed transcribed DNA prior to integration and to G-to-A coding strand mutations of viral genes in the integrated provirus. *APOBEC3* genes are highly evolving and show strong signs of positive selection; the number of *APOBEC3* genes varies from species to species, from 1 gene in mice to 7 genes in primates (1, 10). Human APOBEC3G and 3F were first shown to inhibit HIV-1 lacking the *vif* gene, which encodes a protein expressed at high levels late in infection (11-15). In Vif-deficient-HIV producer cells, APOBEC3 proteins are packaged into progeny virions via interaction with the nucleocapsid protein and viral RNA (16-20).

APOBEC3 proteins also inhibit replication by a number of CDA-independent mechanisms (21). *In vitro* studies have suggested that APOBEC3 proteins inhibit elongation and accumulation of HIV-1 reverse transcription products and we and others have shown that mouse APOBEC3 mostly restricts MLV and mouse mammary tumor virus (MMTV) by inhibiting reverse transcription both *in vivo* and *in vitro* (22-27). Mouse retroviruses are not refractory to APOBEC3-mediated deamination, however, since both *in vitro* and *in vivo* studies using cells and mice transgenic for human APOBEC3G have demonstrated extensive deamination of MLV and MMTV sequences (28-30).

The role of UNG in uracil removal from APOBEC3G-deaminated DNA has been studied in tissue culture cells, with conflicting conclusions (31-34). Here, we tested whether UNG contributed to the repair of APOBEC3G-mediated deamination of replicating MLV *in vivo*, by generating human APOBEC3G (A3G) transgenic mice that lacked the *Ung* as well as the mouse *Apobec3* genes. We found that A3G+, *Ung*-containing mice were more highly infected with MLV than A3G+ *Ung* knockout mice and that proviral DNA from the latter strain had substantially more G-to-A mutations. *In vitro* studies showed that more APOBEC3G-deaminated proviral DNA was integrated into chromosomes in the absence of UNG, suggesting that UNG removal of uracils from unintegrated viral nuclear DNA prevents its integration. These data demonstrate that UNG can counteract the DNA damage inflicted by APOBEC3 deamination.

## Results

We previously reported that transgenic mice expressing human APOBEC3G and deficient in mouse APOBEC3 (APO-/-) were less infected by MLV and that the proviruses found in these mice showed high levels of G-to-A mutations (29). To determine if UNG played a role in the repair of these mutations, we generated UNG-/-APO-/- and A3G^high^UNG-/-APO-/- mice (heterozygous for the A3G^high^ allele) (29). UNG knockout mice are viable and do not display a phenotype other than altered class-switch recombination, but accumulate uracil in their genome (35). Peripheral blood mononuclear cells from A3G^high^ mice express APOBEC3G at levels similar to humans (29). The UNG-/-APO-/-and A3G^high^UNG-/-APO-/-mice were crossed and newborn pups from this cross were infected with MMLV. Newborn APO-/- and A3G^high^APO- /- mice from similar heterozygote crosses were also infected for comparison. At 16 days and 1-month post-infection, MLV titers in the spleens or blood of these mice, respectively, were determined, followed by genotyping for the A3G transgene. Integrated DNA at both time points and viral RNA levels at 16 days post-infection (dpi) were also determined. At 16 days and 1 month post-infection, expression of APOBEC3G reduced *in vivo* infection by ∼2 logs in the spleen and peripheral blood, in both the presence and absence of UNG compared to the non-transgenic APO-/- and UNG-/-APO-/-mice (Fig. 1A, 1B and 1D) (28, 29). Infection levels were higher in the A3G^high^UNG-/-APO-/-mice than in the A3G^high^APO-/- (∼3-fold higher titers at both time points). Splenic viral RNA and DNA levels were also reduced by 1 log at 16 dpi in the A3G^high^UNG-/-APO-/-compared to the A3G^high^APO-/- mice (Fig. 1B and 1C). APO-/- and UNG-/- APO-/- mice showed no significant difference in infection.

**Fig. 1.**
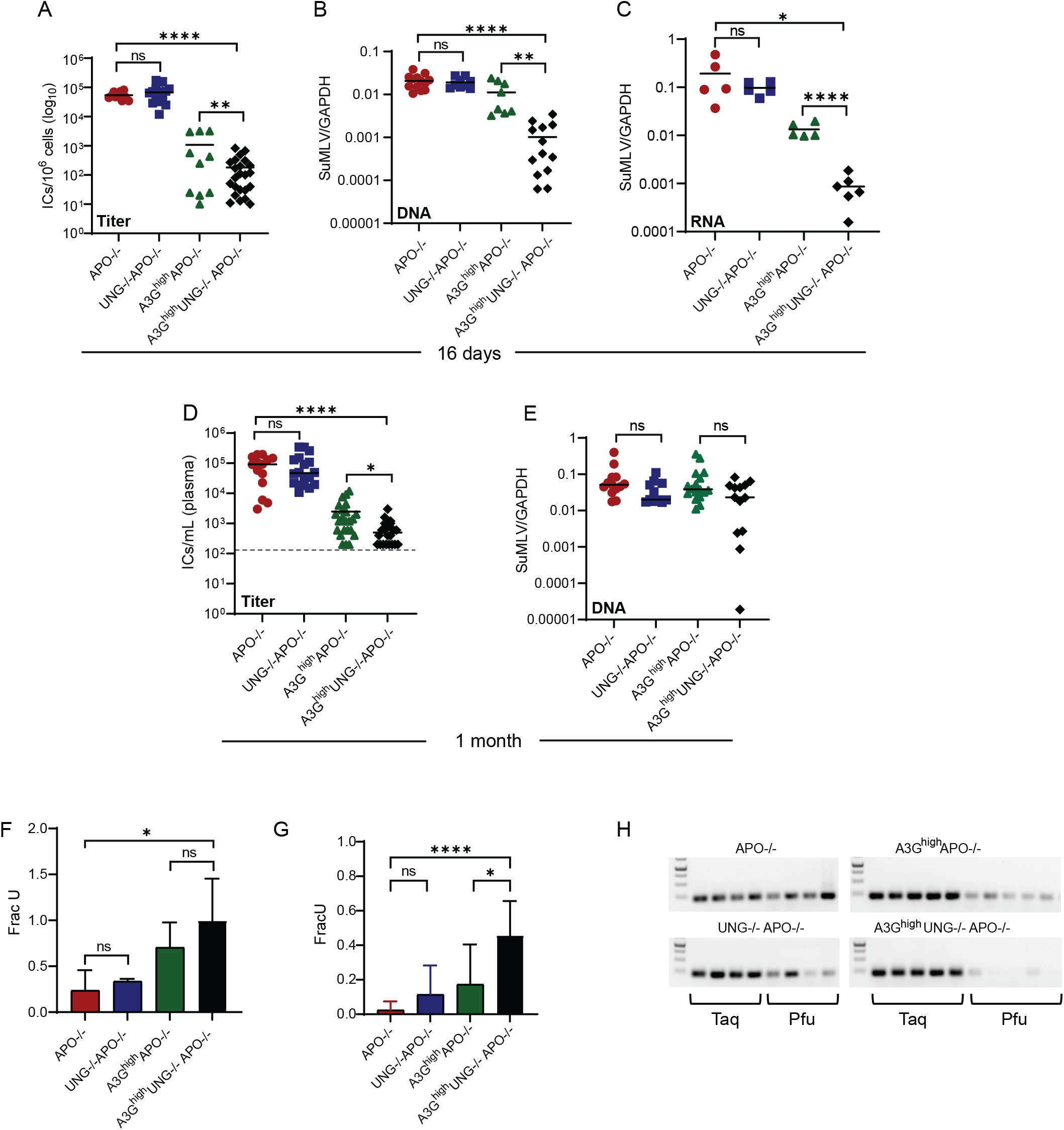
APOBEC3G restrict murine retrovirus infection *in vivo*, in both the presence and absence of UNG. A) Newborn mice of the indicated genotypes were infected with 2x 10^3^ infectious center units (ICs) of MLV, sacrificed at 16 days dpi, and virus titers in spleens were measured. N= 11 APO-/-, 19 UNG-/-APO-/-, 10 A3G^high^APO-/- and 23 A3G^high^UNG-/- APO-/- mice. B) DNA was isolated from spleens at 16 dpi and subjected to qPCR with primers specific to MMLV Env (SuMLV). N= 13 APO-/-, 9 UNG-/-APO-/-, 8 A3G^high^APO-/- and 13 A3G^high^UNG-/- APO-/- mice. C) Viral RNA levels were analyzed by RT-qPCR from spleens at 16 dpi. N= 5 APO-/-, 5 UNG-/-APO-/-, 5 A3G^high^APO-/- and 6 A3G^high^UNG-/- APO-/- mice. D) Mice infected with MMLV were bled at 1-month post-infection, and virus titers in plasma were measured. N= 14 APO-/-, 18 UNG-/-APO-/-, 23 A3G^high^APO-/- and 22 A3G^high^UNG-/- APO-/- mice. E) DNA was isolated from PBMCs at 1-month post-infection and subject to qPCR with SuMLV primers. For A- E, each point represents the data obtained from an individual mouse; the average for each group is shown by a horizontal bar (limit of detection = 200 copies/mL; dashed line). F) Fraction of proviral DNA from spleen at 16 dpi that contain uracil as determined by the Ex-qPCR method, using SuMLV primers specific to the *env* gene. G) Fraction of proviruses from PMBCs at 1-month post-infection that contain uracil as determined by Ex-qPCR. Values represent the mean ± SD from at least 7 different mice. Unpaired two-tailed t tests were used to determine significance. ****, *P* ≤0.0001; **, *P* ≤ 0.006; *, *P* ≤0.05; ns, not significant. H) DNA isolated from the spleens of MLV-infected mice of the indicated genotypes at 16 dpi was amplified with Taq or Pfu polymerase. Each lane is DNA from an individual mouse.

We also examined uracil incorporation in MLV DNA, using 2 different techniques. First, a PCR-based technique developed by the Stiver lab was used to determine the fraction of uracil in integrated DNA (34). At 16 dpi, there was more uracil incorporated in the MLV sequences found in the spleens both UNG^+^ and UNG^-^ A3G^high^APO-/- mice compared to the non-transgenic strains (Fig. 1F). Moreover, the highest levels of uracil were detected in the A3G^high^UNG-/-APO-/-DNA samples. Similar results were seen at 1-month post-infection (Fig. 1G), although the integrated DNA levels were not significantly different (Fig. 1E). We also used a second technique to examine uracil incorporation, that relies on the inability of Pfu polymerase to elongate in the presence of uracil compared to Taq polymerase (36). DNA from the infected spleens of UNG^+^ and UNG^-^ A3G^high^APO-/- mice (16 dpi) amplified more poorly with Pfu polymerase than those from APO-/- and UNG-/-APO-/-mice (Fig. 1H). Moreover, DNA from the A3G^high^UNG-/-APO-/- mice hardly amplified with Pfu polymerase. These data suggest that C-to-U mutations introduced by APOBEC3G into proviruses are not efficiently repaired in the absence of UNG.

### Proviruses in the DNA of A3G^high^UNG KO mice have more G-to-A coding strand mutations

We next subjected DNA isolated from the spleens of individual mice to NextGen sequencing, using primers that spanned the viral genome and that did not amplify endogenous MLV sequences (Fig. 2A). The proviral DNA isolated from the infected spleens of A3G^high^UNG-/-APO-/-mice had almost 2 times more G-to-A mutations than the A3G^high^APO-/- mice and both had mutations at >10-fold higher levels than their non-transgenic counterparts (Fig. 2B). No other types of mutations, including C-to-T mutations indicative of non-coding strand deamination or errors introduced by BER, varied between the different mouse strains (Fig. 2B). The G-to-A mutations were high in the UNG-A3G^high^mice in all regions of the genome compared to the UNG+ A3G^high^ mice (Fig. 2C). Interestingly, in addition to there being a hotspot for APOBEC3G mutations in the 3’ end of the provirus, as has been seen for other retroviruses, there was a second hotspot in the gag gene (red arrows in Fig. 2C). The G-to-A mutations were predominantly found in the APOBEC3G motif GG in the coding strand of both the UNG+ and UNG-A3G^high^ transgenic mice (Fig. 2B and 2D).

**Figure 2.**
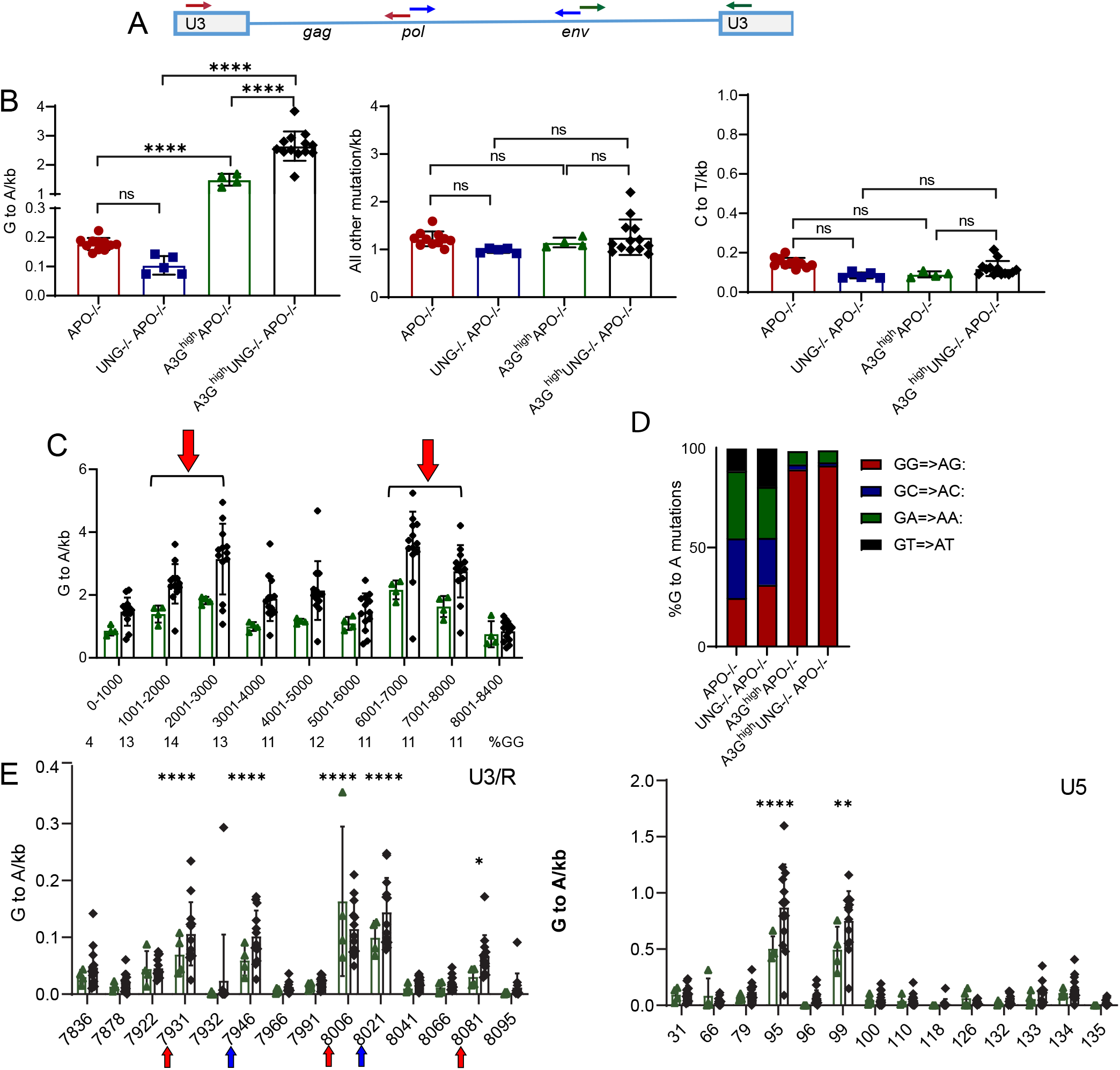
Deamination in the proviral DNA of infected transgenic mice. A) The MLV provirus is shown with the locations of six primers that cover the viral genome. B) DNA was isolated from spleens at 16 dpi and subjected to NextGen sequencing to determine G-to-A mutations, all other mutations and C to T mutations in the proviral DNA. C) G-to-A mutations present across the proviral genome. Red arrows show two hotspots for APOBEC3G mutations in the *gag* (1000 – 3000) and *env* (6000 – 8000) genes. The percent of GG context in each region of the provirus is showed below the x-axis. Each point represents the G-to-A, C-to-T, or all other mutations per kb obtained from an individual mouse. N= 11 APO-/-, 5 UNG-/-APO-/-, 4 A3G^high^APO-/- and 13 A3G^high^UNG-/- APO-/- mice. D) Bar chart showing the percentage of G-to-A mutations in the GG context in proviral DNA. E) G-to-A mutations in the U3/R and U5 regions. Numbering refers to position in viral RNA. Red arrows indicate GREs and blue arrows NFAT1 consensus sequences. Two-way ANOVA with Tukey’s multiple-comparison test was used to determine significance. \*\**P*≤0.001; ****, *P*≤0.0001; ns, not significant.

We also examined G-to-A mutations in the long terminal repeats (LTRs). We found several hotspots in both the U3 and U5 regions (Fig. 2E). Interestingly, the hotspots in U3 occurred in glucocorticoid response elements and binding sites for NFAT1, known to be important for MLV transcription (37). There were two additional hotspots of unknown significance in U5.

These data confirm our previous findings that APOBEC3G mutates MLV and that the absence of UNG leads to even higher G-to-A changes. The mutations found in the MLV-infected A3G^high^UNG-/-APO-/-mice could have a greater effect on both coding regions and virus transcription, thereby decreasing *in vivo* infectivity.

### G-to-A mutations in UNG-/- and UNG+/+ mice are lower in viral RNA than DNA

The proviral DNA isolated from UNG-A3G^high^mice showed substantially more mutations than that from UNG-containing A3G^high^ transgenic mice, and both virus titers and splenic viral RNA levels were reduced. We next examined whether there was a difference in the mutation level in viral RNA isolated from the spleens of mice 16 dpi. As was seen with the viral DNA, RNA from both strains of A3G^high^ transgenic mice had significantly more G-to-A mutations than the nontransgenic strains (Fig. 3A). However, while the G-to-A mutation level was 3-fold higher in DNA vs. RNA for both strains, the level of G-to-A mutations was similar in the viral RNA of the UNG+ and UNG-A3G^high^ transgenic mice, although as was seen for the mutation level in DNA (Fig. 2), there was more variability in the latter strain (Fig. 3B). Both the level of nonsynonymous mutations and stop codons was higher in the proviral DNA of the A3G^high^UNG-/-APO-/-mice than the A3G^high^ APO-/- mice (Fig. 3C). This suggests that only the less heavily mutated proviruses are able to replicate.

**Figure 3.**
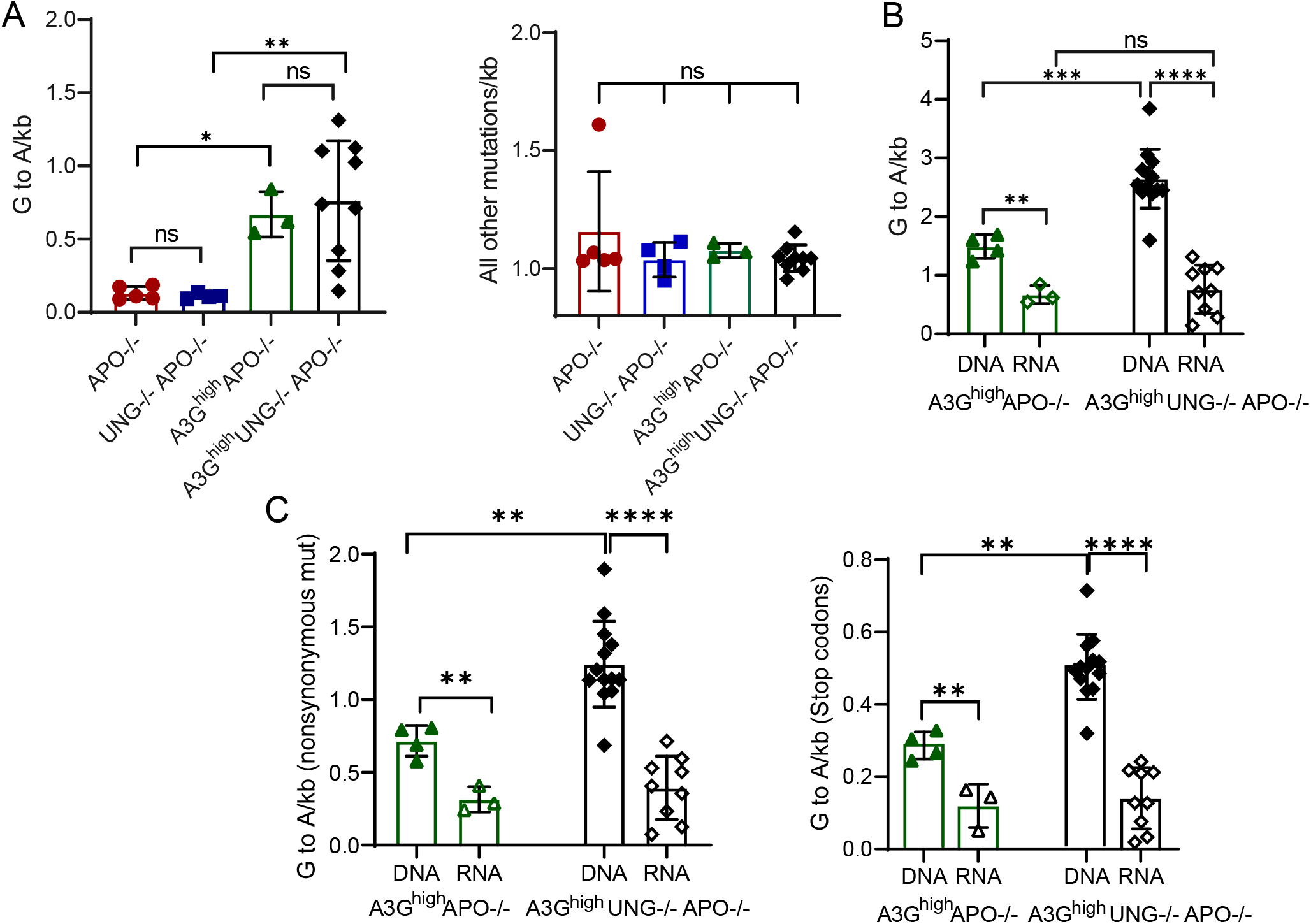
G-to-A mutations in the viral RNA of infected transgenic mice. A) RNA was isolated from spleens at 16 dpi and subjected to NextGen sequencing to determine G-to-A mutations and all other mutations in the viral RNA. B) Comparison of G-to-A mutations between DNA and RNA in A3G^high^UNG-/- APO-/- mice and A3G^high^APO-/- mice are shown. C) G-to-A mutations that cause nonsynonymous mutations and stop codons were analyzed in DNA and RNA. Comparison of the level of nonsynonymous mutations and stop codons between DNA and RNA in A3G^high^UNG-/- APO-/- mice and A3Ghigh APO-/- mice are shown. Each point represents the G-to-A or all other mutations per kb obtained from an individual mouse. N= 5 APO-/-, 4 UNG-/- APO-/-, 3 A3G^high^APO-/- and 9 A3G^high^UNG-/- APO-/- mice. ANOVA with Tukey’s multiple-comparison test was used to determine significance. ****, *P* ≤0.0001; ***, P ≤0.001**, *P* ≤ 0.01; *, *P* < 0.05; ns, not significant.

### Integration levels are higher in UNG-depleted cells

UNG is the major mammalian uracil deglycosylase that removes uracil from genomic DNA (35). The increased mutational burden in the proviral DNA found in A3G^high^ mice that lacked UNG could be due lack of removal of uracil from unintegrated viral DNA or from integrated proviruses. To test at which step uracils are removed, we performed *in vitro* time course assays. First, we generated 293T-MCAT cells, which stably express the MLV receptor mCAT-1, that also expressed APOBEC3G. These cells, as well as 293T-MCAT cells not expressing APOBEC3G, were infected with MLV, and APOBEC3G-containing virus as well virus lacking APOBEC3G was isolated from the supernatants (Fig. 4A). Because APOBEC3G blocks replication, virus stocks were normalized by measurement of virion RNA and by western blot analysis (Fig. 4A; Materials and Methods). Equal amounts (virus RNA equivalents) of APOBEC3G-containing and –lacking viruses were used to infect 293-MCAT cells which were treated with UNG or control siRNAs, and at 2, 4, 6, 8, and 24 hours post-infection (hpi), the cell extracts were fractionated into cytoplasmic, nuclear soluble and insoluble fractions (Fig. 4B). UNG knockdown was confirmed by RT-qPCR (Fig. 4A). DNA was isolated from each of the fractions and subjected to qPCR to measure viral DNA levels, as well as to analysis of uracil content.

**Figure 4.**
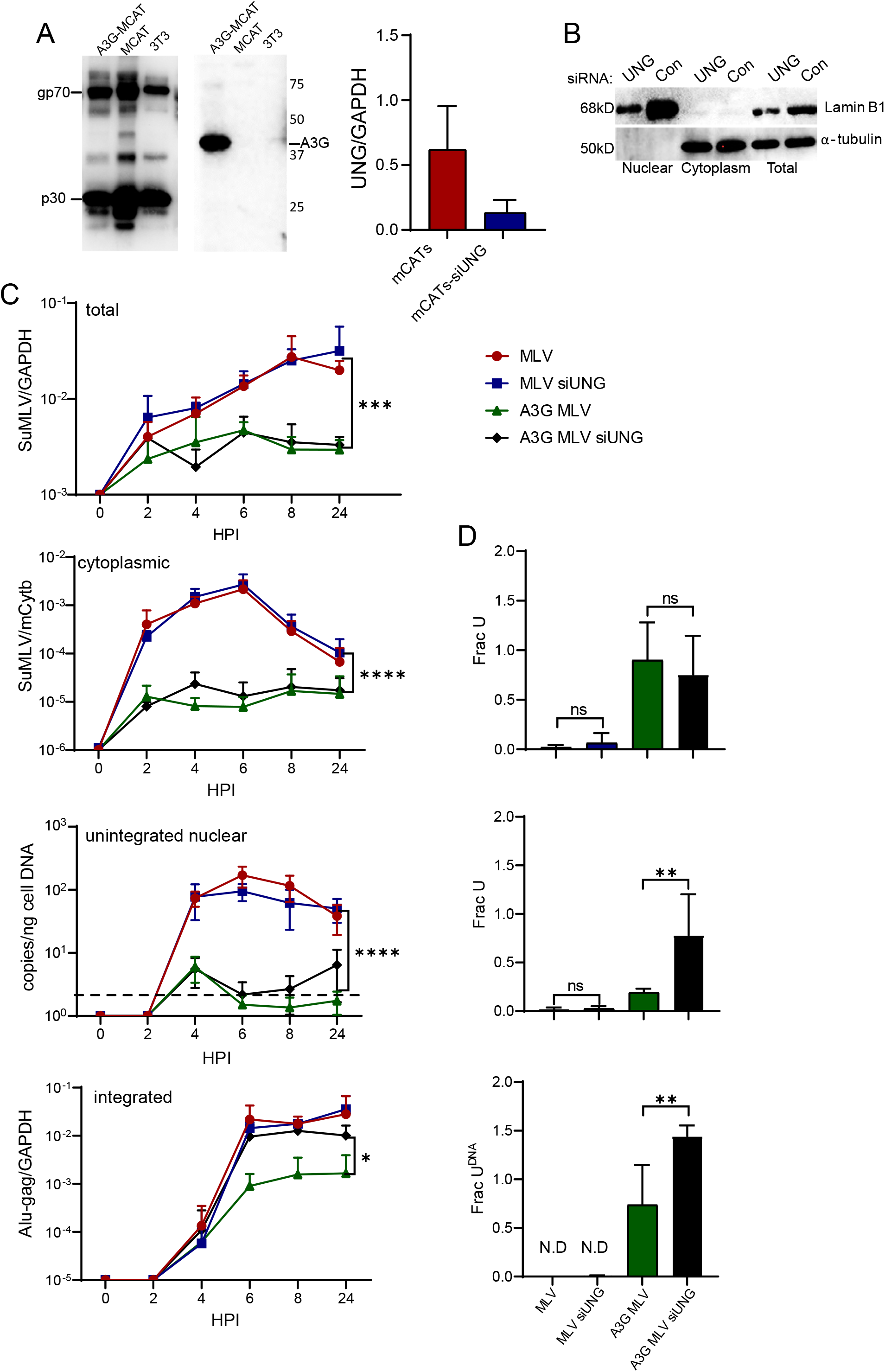
UNG removes uracils from unintegrated nuclear DNA and avoids integration. A) Left panel: western blots showing MLV viruses from the different cell lines used, and APOBEC3G expression in MLV virus from A3G MCAT cells; the panel on the right shows the UNG knockdown. B) Fractionation and western blots of the cells used in part C. Laminin B1 and α- tubulin were used as markers for the nucleus and cytoplasm, respectively. C) Equal amounts of MLV were used to infect 293-MCAT cells which were transfected with UNG or control siRNAs, at 2, 4, 6, 8, and 24 hpi, the cell extracts were fractionated into total, cytoplasmic, unintegrated nuclear and integrated fractions. DNA was isolated from each of the fractions and subjected to qPCR to measure viral DNA levels using SuMLV primers for total, cytoplasmic and unintegrated nuclear fractions. Integrated MLV DNA quantification was performed by real-time Alu-gag qPCR. Values were normalized to GAPDH for total and nuclear fractions, to mCytb for cytoplasmic fraction and to total DNA for the unintegrated nuclear fraction. Each point shows the averages ± SD of 3 different experiments. Two-way analysis of variance (ANOVA) with Tukey’s or Šídák’s multiple-comparison test was used to determine significance. ****, *P* <0.0001; ***, *P* <0.001; * *P* < 0.05. D) Ex-qPCR method with SuMLV primers was used to determine the fraction of viruses containing uracils in the cytoplasmic, unintegrated and high molecular weight nuclear fractions at 6 hpi. Bars show the average ± SD of 4 different experiments. One-way ANOVA with Tukey’s multiple-comparison test was used to determine significance. **, *P* ≤ 0.01; ns, not significant.

Virus reverse transcription was diminished in cells infected with APOBEC3G-containing virus, irrespective of the expression of UNG. This was true for all unintegrated forms (nuclear and cytoplasmic) of viral reverse transcripts (Fig. 4C). However, while proviral integration levels remained low in cells infected with the APOBEC3G-containing virus and expressing UNG, in the UNG knockdown cells, levels of integrated viral DNA were almost at the level as in cells infected with virus lacking APOBEC3G (Fig. 4C, integrated). When uracil incorporation into the viral DNA from different fractions was determined, we found that uracil levels were higher in DNA isolated from all fractions of cells infected with APOBEC3G-containing virions (Fig. 4D). Moreover, while uracil levels in cytoplasmic viral DNA in the UNG-expressing and -depleted cells infected with APOBEC3G-containing virus were similar, the levels in unintegrated nuclear and integrated proviral DNA from the UNG-depleted cells were higher than that from UNG-expressing cells. (Fig. 4D). Taken together, these data suggest that 1) UNG removes uracils from unintegrated viral DNA in the nucleus, and this nicked DNA integrates less efficiently than uracil-containing, intact viral DNA; and 2) the uracil found in proviruses made in the absence of UNG causes increased G-to-A mutations.

## Discussion

Previous studies have disagreed as to whether UNG is involved in the repair of APOBEC3-mediated cytidine deamination of retroviral DNA (reviewed in ref. (38)). However, many of these studies were done with over-expressed APOBEC3 or UNG proteins, and used short-term replication assays to assess the effects of UNG. Here, we show using an *in vivo* system, in which virus undergoes multiple rounds of replication, that UNG plays a role in removing the uracils introduced by APOBEC3G-mediated cytidine deamination into MLV proviruses. As we showed previously, an APOBEC3G transgene expressed at levels similar to that seen in humans, introduces “catastrophic” G-to-A mutations into the coding strand of MLV-infected mice, reducing in vivo infection by several logs. In A3G transgenic mice that also lack *Ung*, the G-to-A mutation rate was increased to even higher levels, which resulted in lower levels of infection. Thus, UNG could be characterized as a pro-viral factor that aids in the repair of mutations introduced into the viral genome by the APOBEC3 cytidine deaminases. That lack of UNG did not cause even higher rates of mutation and greater effects on infection is likely due to the other BER enzymes that repair uracil in DNA, such as selective monofunctional uracil-DNA glycosylase (SMUG1), thymidine DNA glycosylase (TDG) and methyl CpG binding domain 4 (MBD4) (35).

In *in vitro* studies, incorporation of APOBEC3G into MLV particles reduced cytoplasmic and unintegrated reverse transcripts, as well as integrated DNA, independent of UNG expression compared to virions lacking APOBEC3G. This is likely because APOBEC3G, in addition to deaminating newly synthesized viral DNA, can block reverse transcription (9). When APOBEC3G-containing MLV was used to infect tissue culture cells in which UNG levels were reduced by siRNA, the level of unintegrated nuclear DNA was similar in the UNG-expressing and –negative cells infected with APOBEC3G-containing MLV. In contrast, we found that integration of proviral DNA was increased in UNG-depleted cells relative to cells expressing UNG. Additionally, the level of uracil incorporated in nuclear unintegrated viral and proviral DNA was higher in the UNG-deficient cells compared to the UNG-expressing cells. This suggests that when UNG acts on unintegrated viral DNA, the cleavage sites are not repaired by the BER machinery, likely leading to nicked DNA that does not efficiently integrate. In contrast, proviruses containing uracil would be cleaved by UNG after integration, and repaired using the cellular BER machinery; this would not occur in UNG-deficient cells and could explain the higher G-to-A mutation rate in the A3G^high^UNG-/- mice compared to the A3G^high^ mice. The repair of uracil in integrated proviruses by UNG also explains why virus replication levels were higher in A3G^high^ APO-/- than A3G^high^ UNG-/-APO-/ mice. Although BER is known to be error-prone, we did not see evidence of increased mutations other than G-to-A, suggesting that instead DNA polymerase recognized uracils as thymidines in the integrated proviruses during DNA replication.

HIV’s replication complexes, consisting of viral capsid, reverse transcriptase, integrase and nucleic acid, can enter the nucleus through interaction with the nuclear pore, and as a result, HIV can infect quiescent cells (39). MLV, in contrast, requires cell division and nuclear membrane breakdown for complex entry because it lacks viral proteins that interact with the nuclear pore complex; it thus can only efficiently infect cycling cells (39). Recent studies have suggested that HIV reverse transcription largely occurs in the nucleus (40-43). Whether this is also the case for gammaretroviruses is not known. However, cytoplasmic viral DNA isolated from cells infected with APOBEC3G-containing virus had significant levels of uracil, suggesting that at least some reverse transcription occurs prior to association of the reverse transcription complex (RTC) with the nucleus (Fig. 4D). However, the level of uracil in cytoplasmic DNA did not differ in UNG-containing and –depleted cells, but did in the nuclear fractions. Thus, UNG, which is a nuclear enzyme, is likely removing uracils in the nucleus. Although UNG can remove uracils from double-stranded DNA, its activity is higher on single-stranded DNA, such as that occurs at replication foci or during reverse transcription (35). If some reverse transcription and APOBEC3G-mediated deamination occurs in the nucleus or during cell division, then nuclear UNG could cause nicks in unintegrated viral DNA through base excision. Further studies are required to elucidate how and where MLV reverse transcription, APOBEC3G-deamination and UNG excision occur.

## Materials and Methods

### Ethics statement

All mice were housed according to the policy of the Animal Care Committee of the University of Illinois at Chicago, and all studies were performed in accordance with the recommendations in the Guide for the Care and Use of Laboratory Animals of the National Institutes of Health. The experiments performed with mice in this study were approved by the committee (UIC ACC protocol #18-168).

### Mice

A3G^high^APO -/- mice and APO-/- mice were previously described (23, 29). UNG-/- were a generous gift from Amy Kenter (6). Conditions for genotyping the A3G transgene, as well as the mouse *Apobec3* gene, were reported previously (23, 29). Knockout of the *Ung* gene was verified using the following primers: (UNGKO F primer 5’-GCCGGTCTTGTCGATCAGGATGATC-3’ and UNGKO R primer 5’-CAGTGCCTATAACTTCAGCTCC-3’).

### Cell culture

NIH3T3 cells were cultured in Dulbecco’s modified Eagle’s medium (DMEM) supplemented with 10% fetal bovine serum (FBS), L-glutamine, and penicillin/ streptomycin. 293T/mCAT-1 cells were a gift from Lorraine Albritton (44). 293T/A3G/mCAT-1 cells expressing human APOBEC3G were generated by co-transfecting A3G expression and puromycin-resistance plasmids. The 293T/mCAT-1 and 293T/A3G/mCAT-1 cells were cultured in Dulbecco’s modified Eagle’s medium (DMEM) supplemented with 8% donor bovine serum (DBS), L-glutamine, penicillin/ streptomycin containing G418 (Goldbio) or G418 plus puromycin (Gibco), respectively.

### Virus isolation

MMLV was isolated from the supernatants of stably infected NIH3T3 cells (cells in which infection is allowed to spread to 100% of the culture and maintained in this state thereafter), as previously described (25, 45). Virus was also isolated from MLV-infected 293T/mCAT-1 and 293T/A3G/mCAT-1 cells. Supernatants were passed through a 0.45-µm filter, treated with 20 U/ml DNase I (Sigma) at 37°C for 30 min and centrifuged through a 30% sucrose cushion, as previously described. After resuspension, titers of MLV were determined on NIH3T3 cells (see “Virus titers,” below).

Viruses were subjected to reverse transcriptase quantitative PCR (RT-qPCR), and the number of viruses was estimated by standard curve analysis from the amount of virus-specific RNA, using primers located in the env gene (MMLV F primer, 5’-CCTACTACGAAGGGGTTG-3’; MMLV R primer, 5’-CACATGGTACCTGTAGGGGC-3’). Equal amounts of virus, normalized by RNA levels, were also analyzed by Western blots (Fig. 4A).

### In vivo infections

One-to-2-day-old mice were infected by intraperitoneal injection of 2 × 10^3^ ICs of MMLV and spleens were harvested at 16 days dpi, as previously described (17). Mice were anesthetized and blood was obtained via retro-orbital bleed. Plasma and peripheral blood mononuclear cells were collected with heparinized Natelson tubes (Fisher Scientific) into 8mM EDTA in PBS. Plasma samples were serially diluted to titer virus. For cellular DNA isolation, red blood cells were lysed with ACK lysis buffer (150mM NH4Cl, 1 M KHCO3, 0.1mM EDTA, pH 7.4) and cells were washed twice with PBS and finally diluted in 200 uL of PBS. Samples were stored at -20 °C prior DNA isolation.

### Virus titers

MMLV infection levels in the spleens and peripheral blood of the infected mice or the supernatants of infected 293T/mCAT-1 and 293T/A3G/mCAT-1 cells were determined by infectious center (IC) assays using a focal immunofluorescence assay, as previously described (37). Briefly, NIH3T3 cells were infected with 10-fold serial dilutions of splenocytes or virus, respectively. At 4 dpi, the plates were stained a monoclonal antibody (538) that recognizes the Env protein. After staining with fluorescein-conjugated secondary antibody, the colonies of green cells were quantified by automated counting using a Keyence fluorescence microscope. Viral titers (ICs) were calculated from the numbers of fluorescent colonies corrected for the dilution factors of the viral stocks in each plate.

### Deep Sequencing of nearly full-length MMLV genomic DNA and RNA

DNA from the spleens of MLV-infected APO-/- A3G^high^, UNG-/-APO-/- A3G^high^ mice and APO-/-, UNG-/-APO-/-control mice was isolated using the DNeasy Blood & Tissue Kit (Qiagen). RNA was also isolated from MMLV-infected splenocytes of the mice using Trizol reagent (Ambion), and cDNA was reverse transcribed using AccuScript High Fidelity First-Strand cDNA Synthesis kit (Agilent Technologies). Three MMLV fragments that covered most of the proviral genome were amplified from DNA and RNA using the primers described in Table I (Fig. 2A; Table 1). Briefly, the three amplicons were purified (Agencourt AMPure XP) and quantified (Nanodrop) prior to using the Celero™ DNA-Seq Library Preparation Kit (NuGEN) to construct libraries. These libraries were analyzed using an Agilent Tapestation 4200 for size and concentration (Agilent Technologies). Libraries were then pooled based on nM concentration and the resulting pool prepared for sequencing by measuring concentration by Qubit 4 (Life Technologies). The pooled libraries were run on an Illumina MiniSeq instrument at 2 × 150bp using MiniSeq Reagent MO Kit, (300 cycles) (#FC-420-1004 Illumina Inc).

**Table 1:**
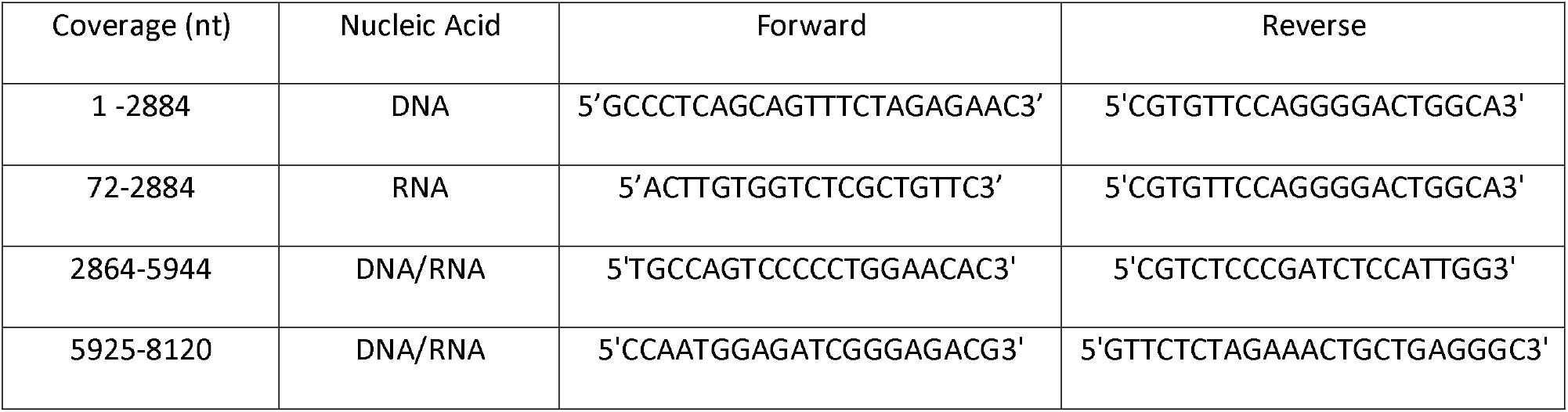
Primers used to amplify MMLV proviral DNA and viral RNA genome.

### Sequence analysis

Raw reads were mapped to the Moloney murine leukemia virus (J02255) using BWA MEM(46) (total mapped reads average: 1.3 × 10^5^). PCR duplicates were removed using Picard MarkDuplicates (47), and indel realignment was performed using IndelRealigner from GATK(48). Nucleotide counts per position were generated at each position in the reference using bam-readcount (Bam-Readcount: Generate Metrics at Single Nucleotide Positions., n.d.) and the effect of substitutions on the translated protein sequence were assessed for the open read frames in the virus: positions 621-2237 (Gag polyprotein pr65), positions 2238-5834 (Pol polyprotein), and 5777-7774 (Env polyprotein). Distributions of both single nucleotide conversions and dinucleotide conversions were compiled over all positions in the genome, in particular G->A and C->T conversions for single nucleotides, and GG->AG, GC->AC, GA->AA, and GT->AT conversions for dinucleotides. These conversion frequencies were also averaged over 1kb bins across the reference sequence. Differential statistics of conversion frequencies between sample groups were tested using the Wilcox test in R. G to A/kb, C to T/kb and all mutations/kb calculations were made counting total number of G to A, C to T or the rest of mutations and dividing these numbers between kb reads.

### RNAi

For the depletion of UNG in human cells, siRNA from Ambion (catalog no. 4390824) was used. Briefly, 2293T/mCAT-1 and 293T/A3G/mCAT-1 were transfected using the reverse-transfection method of Lipofectamine RNAi MAX reagent (Invitrogen). siRNA depletion was carried out for 48 h. RNA was isolated using the RNeasy minikit (Qiagen). RT-qPCR was performed using the GoTaq® 1-Step RT-qPCR System (Promega). Knockdowns were verified using the primers: 5’CTCATAAGGAGCGAGGCTGG3’ and 5’GTACATGGTGCCGCTTCCTA3’.

### In vitro infections to determine reverse transcription early events

293T/mCAT-1 and 293T/A3G/mCAT-1 cells were seeded at 1 ⍰ × ⍰ 10^5^ cells per 0.5⍰ ml of medium in a 24-well format. Virus (genome equivalent of a MOI of 1) was added in the presence of 8 μg/ml polybrene (Sigma Aldrich) and the cells were incubated on ice for 1⍰ h to allow virus binding. Cells were washed in cold phosphate-buffered saline, 0.5 ml of DMEM was added, and incubated at 37 °C for 0-6 h, as indicated in the figures. At each harvest time point, the cells were fractionated by the modify rapid, efficient, and practical (REAP) method as previously described (49). Total, integrated and cytoplasmic DNA was purified from the REAP fractions using DNeasy kits (Qiagen). The purity of the fractions was determined by western blotting with antibodies to β-tubulin (cytoplasmic fraction) (GeneTex) and laminin B1 (nuclear fraction) (Cell signaling Technology). Unintegrated nuclear DNA was isolated using the Hirt DNA isolation method, appropriate for extraction of low molecular weight DNA (50). Briefly, Hirt buffer (0.09M Tris pH7.6, 0.01M EDTA, 0.6% SDS) was added to the REAP nuclear fraction, and incubated for 10 minutes. After, ¼ volume of 5.0M NaCl was added and mixed gently. The lysis mixture was incubated at 4°C overnight. The mixture was centrifugated at 13,000 rpm for 15 min at 4°C, and then supernatant was carefully removed, mixed with Proteinase K (0.1 mg/mL), and incubated at 56°C for 2 hours, followed by phenol-chloroform extraction and ethanol precipitation. The pellet was diluted in Phosphate-buffered saline (PBS) to isolated the integrated DNA. The DNA from the different fractions was subjected to real-time qPCR.

### Real-time qPCR

qPCRs were performed with MLV SuMLV primers using a Power SYBR green PCR kit (Promega) and the QuantStudio 5 Real-Time PCR System (Applied Biosystems). DNA quantifications were normalized to glyceraldehyde-3-phosphate dehydrogenase (GAPDH), or to the mitochondrial gene for cytochrome b (mtCytb) in the cytoplasmic fraction. The amplification conditions were 50°C for 2 min, 95°C for 10 min and 40 cycles of 95°C for 15 s, and 60°C for 1 min. The efficiency of amplification was determined for each primer pair by generating a standard curve with 10-fold serial dilutions of a known concentration of DNA. For each primer pair, a no-template control was included, and each sample was run in triplicate. Levels of integrated MLV were determined by Alu-gag nested PCR (45). Briefly, 50 ng of total DNA was used to perform a PCR using a forward primer that targeted genomic *alu* sequences randomly located near integrated proviruses, and MLV-specific gag reverse primer. The PCR product was diluted 10-fold and 2.4 μl was used as input for the second qPCR reaction, which was performed using MLV LRT primers. The qPCR was normalized to GAPDH using the same amount of input total DNA sample to measure integrated MLV. Copies of unintegrated DNA were determined by qPCR with SuMLV primers and normalized to ng of total DNA in the sample.

### Uracil content of viral DNA

Excision-qPCR was used to determine uracil-containing fraction of viral DNA as described (51) with some modifications. The sample was split into two equal portions and one portion is treated with UDG. Briefly, 0.125 units of UDG (NEB) was added into the Promega qPCR master mix to excise uracils from viral DNA. The qPCR thermocycler reaction was modified to include the UDG reaction time and heat-cleavage of the resulting abasic sites. Thermocycler program we used for this reaction was: 37 °C for 30 min (UDG reaction), 95 °C for 5 min (abasic site cleavage) and 40 cycles of denaturation at 95 °C for 10 sec and annealing and extension at 60 °C for 30 sec. SuMLV primers were used in cytoplasmic, nuclear unintegrated and integrated fraction, to amplify viral DNA. Primers targeting GAPDH or mCytb were used to calculate Frac UDNA using the ΔΔC_t_ method in the nuclear or cytoplasmic fractions, respectively.

The Taq/Pfu PCR method was also used to examine DNA uracil content, as described (36). DNA from spleen at 16 dpi were used as templates for Taq and Pfu amplification with SuMLV primers.

### Statistical analysis and data deposition

Data shown are the averages of at least 3 independent experiments, or as otherwise indicated in the figure legends. Statistical analysis was performed using GraphPad Prism 9.0.2 software. Tests used to determine significance are indicated in the figure legends. Raw data for all figures are deposited in a Mendeley dataset at doi: 10.17632/jmpdfkvd2j.1.

## ACKNOWLEDGEMENTS

We thank David Ryan for assistance with mouse breeding, Alexya Aguilera for providing virus and helpful suggestions and Amy Kenter for providing the UNG KO mice. This study was supported by National Institute for Allergy and Infectious Disease (R01AI 085015 to SRR).

